# Targeting neuronal activity-regulated neuroligin-3 dependency for high-grade glioma therapy

**DOI:** 10.1101/153122

**Authors:** Humsa S. Venkatesh, Lydia T. Tam, Pamelyn J. Woo, Surya Nagaraja, Shawn M. Gillespe, James Lennon, Jing Ni, Damien Y. Duveau, Patrick J. Morris, Jean J. Zhao, Craig J. Thomas, Michelle Monje

## Abstract

Neuronal activity promotes high-grade glioma (HGG) growth. An important mechanism mediating this neural regulation of brain cancer is activity-dependent cleavage and secretion of the synaptic molecule and glioma mitogen neuroligin-3 (Nlgn3), but the therapeutic potential of targeting Nlgn3 in glioma remains to be defined. We demonstrate a striking dependence of HGG growth on microenvironmental Nlgn3 and determine a targetable mechanism of secretion. Patient-derived orthotopic xenografts of pediatric glioblastoma, diffuse intrinsic pontine glioma and adult glioblastoma fail to grow in Nlgn3 knockout mice. Glioma exposure to Nlgn3 results in numerous signaling consequences, including early focal adhesion kinase activation upstream of PI3K-mTOR. Nlgn3 is cleaved from both neurons and oligodendrocyte precursor cells via the ADAM10 sheddase. Administration of ADAM10 inhibitors robustly blocks HGG xenograft growth. This work defines the therapeutic potential of and a promising strategy for targeting Nlgn3 secretion in the glioma microenvironment, which could prove transformative for treatment of HGG.

---

High-grade gliomas (HGG) are a devastating group of tumors, representing the leading cause of brain cancer-related death in both children and adults. Therapies aimed at mechanisms intrinsic to the glioma cell have translated to only limited success, and it is increasingly clear that effective therapeutic strategies will need to also target elements of the glioma microenvironment that promote glioma survival and proliferation. We have recently demonstrated that neuronal activity robustly promotes the growth of a range of molecularly and clinically distinct HGG types, including adult glioblastoma (GBM), anaplastic oligodendroglioma, pediatric GBM, and diffuse intrinsic pontine glioma (DIPG)^1^. Mechanisms mediating this growth-promoting effect of neuronal activity on HGG include activity-regulated secretion of the glioma mitogens brain-derived neurotrophic factor (Bdnf) and neuroligin-3 (Nlgn3) into the tumor microenvironment^1^. Nlgn3 is a post-synaptic adhesion molecule present chiefly at excitatory synapses^2,3^ and expressed by both neurons and by glial cells^4^. Similar to activity-regulated cleavage of neuroligin-1 (Nlgn1)^5,6^, we found Nlgn3 to be cleaved at the c-terminal transmembrane domain, resulting in shedding of the large N-terminal ectodomain^1^. The released Nlgn3 acts on glioma cells to drive proliferation through the PI3K-mTOR pathway, and also promotes a feed-forward increase in glioma NLGN3 expression^1^. Blocking Nlgn3 release into the tumor microenvironment may represent a promising therapeutic opportunity, but the proteolytic mechanism of Nlgn3 cleavage, cell type(s) from which activity-regulated shedding occurs and the relative promise of Nlgn3 as a therapeutic target remain to be clarified.

## Neuroligin-3 necessity to *in vivo* glioma growth

Multiple cell-intrinsic and microenvironmental factors could promote glioma growth, and Nlgn3 is only one such factor. To test the relative necessity of microenvironmental Nlgn3 to glioma growth *in vivo*, we xenografted patient-derived HGG cells expressing GFP-luciferase into *Nlgn3* knockout mice^7^, back-bred onto an immunodeficient background strain to enable xenografting (*Nlgn3*^y/-^; NSG). *Nlgn3* knockout mice are healthy and nearly normal neurologically, exhibiting only subtle deficits in behavioral and electrophysiological assessments^8–10^. A patient-derived culture of a pediatric glioblastoma of the frontal lobe expressing GFP and luciferase (SU-pcGBM2-GFP-luc) was xenografted to the frontal cortex at postnatal day 35 (P35) and monitored using *in vivo* bioluminescent (IVIS) imaging over the course of 6 months (Fig. 1a). Initial engraftment was equivalent in both genotypes (*Nlgn3*^y/-^;NSG and *Nlgn3*^y/+^;NSG; Extended Data Fig. 1). A striking stagnation of glioma growth was evident in the *Nlgn3* knockout animals (at 3 months, tumor burden had increased by ~5-10 fold in *Nlgn3*^y/+^;NSG mice yet remained unchanged in *Nlgn3*^y/-^;NSG mice, n=11 WT, n=14 *Nlgn3* KO, P<0.001) up to six months following xenotransplantation (Fig 1a-f and Extended Data Fig. 2). By four and half months, a subset of tumors circumvents this apparent Nlgn3 dependency and began to exhibit growth (Fig. 1e, f, Extended Data Fig. 2). Inhibition of growth in the Nlgn3^y/-^ mice was demonstrated by IVIS imaging (Fig. 1a, c-f) and confirmed histologically (Fig. 1b)

**Figure 1:**
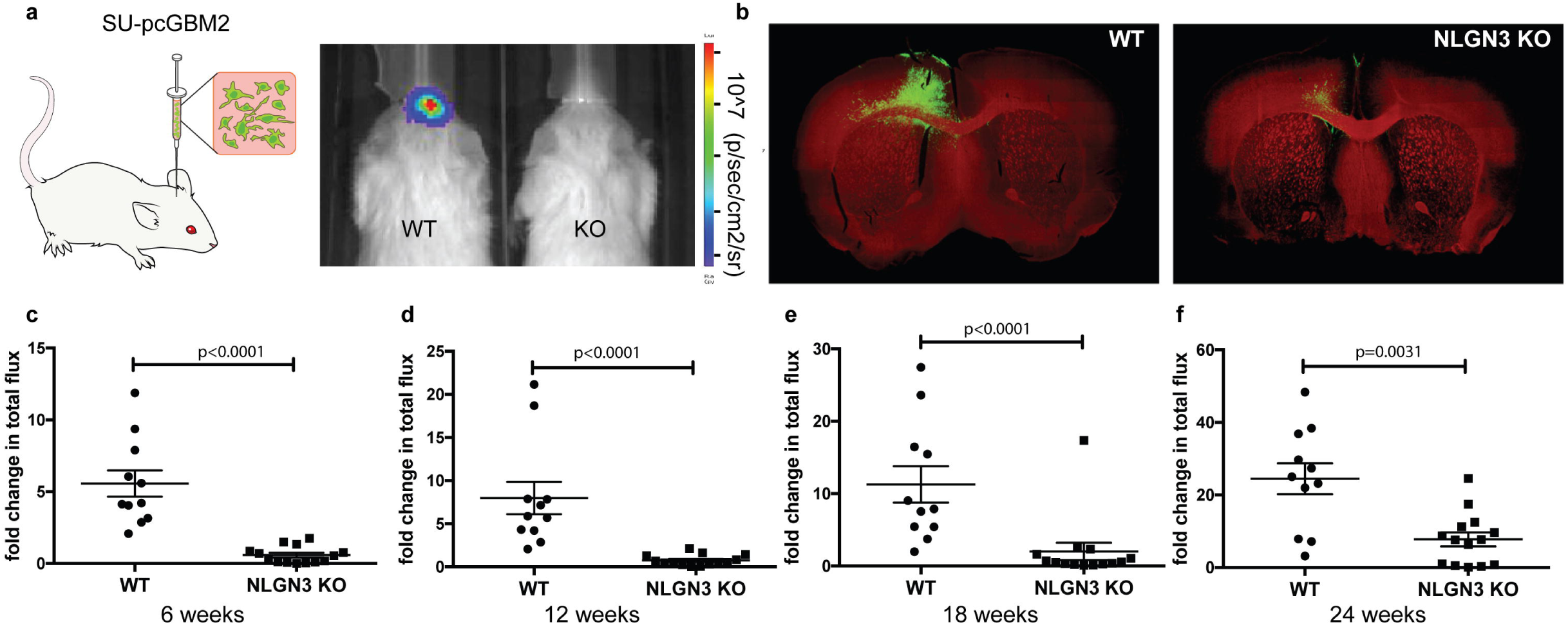
Microenvironmental Nlgn3 is necessary for pediatric GBM growth. **a,** Left: Schematic representation of GFP and luciferase-labeled patient-derived pediatric glioblastoma cells (SU-pcGBM2) orthotopically xenografted into the premotor cortex of immunodeficient WT or *Nlgn3* KO mice Right: Representative IVIS images in WT or *Nlgn3* KO mice at 3 months. Heat map represents degree of photon emission. **b**, Representative confocal micrographs illustrating tumor burden at 6 months; GFP+ tumor cells (green) and myelin basic protein (MBP, red). **c-f**, Fold change in total photon flux of SU-pcGBM2 xenografts in identically manipulated WT;NSG (n=11) and *Nlgn3*^y/-^;NSG (n=14) mice shown at 6, 12, 18, and 24 weeks post-xenograft. Experiment was replicated in five independent cohorts of mice and data are shown combined. Each dot represents one mouse. P values indicated on graphs, Mann-Whitney test. Data shown as mean +/- s.e.m.

The degree of growth inhibition observed in the Nlgn3-deficient mouse brain was unexpected, particularly given our previous work demonstrating that in addition to Nlgn3, Bdnf also contributes to glioma proliferation following exposure to activity-regulated factors present in conditioned medium (CM) from active brain slices^1^. We therefore revisited the necessity of Nlgn3 to glioma cell proliferation in response to active CM *in vitro* to confirm our previous findings regarding a role for other activity-regulated factors. We generated acute cortical slices from optogenetically controllable (Thy1::ChR2), *Nlgn3* knockout mice to enable optogenetic stimulation in the presence or absence of Nlgn3 *in situ*. Acute cortical slices from Nlgn^y/+^;Thy1::ChR2 or Nlgn^y/-^;Thy1::ChR2 mice were optogenetically stimulated *in situ*, and the surrounding conditioned medium (CM) was placed onto patient-derived HGG cultures (Extended Data Fig. 3). Consistent with our previous findings, CM from *Nlgn3*^y/+^;Thy1::ChR2 slices significantly increased the proliferation of HGG cells compared to baseline media, while CM from Nlgn3^y/-^;Thy1::ChR2 elicited a substantially smaller but still significant increase in proliferation (Extended Data Fig. 3). These data replicate the degree of differential proliferation observed when Nlgn3 was sequestered from active slice conditioned medium, with residual proliferation-inducing capacity of the CM accounted for by activity-regulated Bdnf^1^. Taken together, these findings indicate that glioma growth is more dependent on Nlgn3 *in vivo* than would have been predicted from these *in situ*/*in vitro* experiments, at least during early phases of glioma growth. The role that other activity-regulated glioma mitogens such as Bdnf may play at later stages of growth or in the observed ability of some tumors to circumvent dependency on Nlgn3 remains to be determined.

The nearly normal neurological function of *Nlgn3* knockout mice is thought to be due to compensatory expression of other neuroligins^7,11^ such as Nlgn1, a molecule that is also cleaved and secreted in activity-dependent fashion^5,6^. Questioning the apparent lack of compensatory effects of Nlgn1 on glioma growth in the *Nlgn3* knockout mouse, we tested the effects of recombinant Nlgn1 exposure on glioma cell proliferation. Unlike Nlgn3^1^, we did not find a proliferation-inducing effect of Nlgn1 on HGG proliferation *in vitro* (Extended Data Fig. 4). Similarly, we previously demonstrated that neuroligin-2 (Nlgn2) does not promote glioma proliferation^1^. Thus, compensatory expression of other neuroligins would not be expected to influence glioma growth, supporting a unique role for Nlgn3 in glioma pathobiology.

To determine if Nlgn3 plays a similarly important role in the growth of additional types of malignant glioma, patient-derived xenografts of DIPG and adult glioblastoma were tested in the Nlgn3-deficient brain. A patient-derived culture of DIPG (SU-DIPGVI-GFP-luc) was xenografted to the pons at p35 and monitored using *in vivo* bioluminescent imaging, and a similar stagnation of growth was seen in this pontine DIPG xenograft (Fig. 2a-c). At 6 weeks, tumor burden had increased by ~15 fold in *Nlgn3*^y/+^;NSG mice yet remained unchanged in *Nlgn3*^y/-^;NSG mice as determined by *in vivo* bioluminescent imaging (Fig. 2c) and histological analysis (Fig. 2b). A second patient-derived culture of DIPG (SU-DIPGXIII-GFP-luc) isolated from a DIPG frontal lobe metastasis was xenografted to the frontal cortex and similarly displayed growth inhibition in the Nlgn3-deficient brain (Fig. 2d), indicating that DIPG exhibits a dependency on microenvironmental Nlgn3 in both cortical and pontine locations. Similarly, a patient-derived adult glioblastoma culture (SU-GBM035-GFP-Luc) xenografted to the frontal cortex of *Nlgn3*^y/-^;NSG mice exhibit a striking stagnation of growth compared to GBM cells xenografted to the *Nlgn3*^y/+^;NSG mouse brain (Fig. 2e). In contrast, frontal cortex xenografts of a patient-derived HER2^+^ breast cancer brain metastasis (DF-BM354-Luc)^12^ did not exhibit any difference in growth in the presence or absence of Nlgn3 in the brain microenvironment (Fig. 2f, g). These results indicate a conserved dependency on Nlgn3 across molecularly and clinically distinct types of HGG and underscore the importance of microenvironmental Nlgn3 to glioma growth.

**Figure 2:**
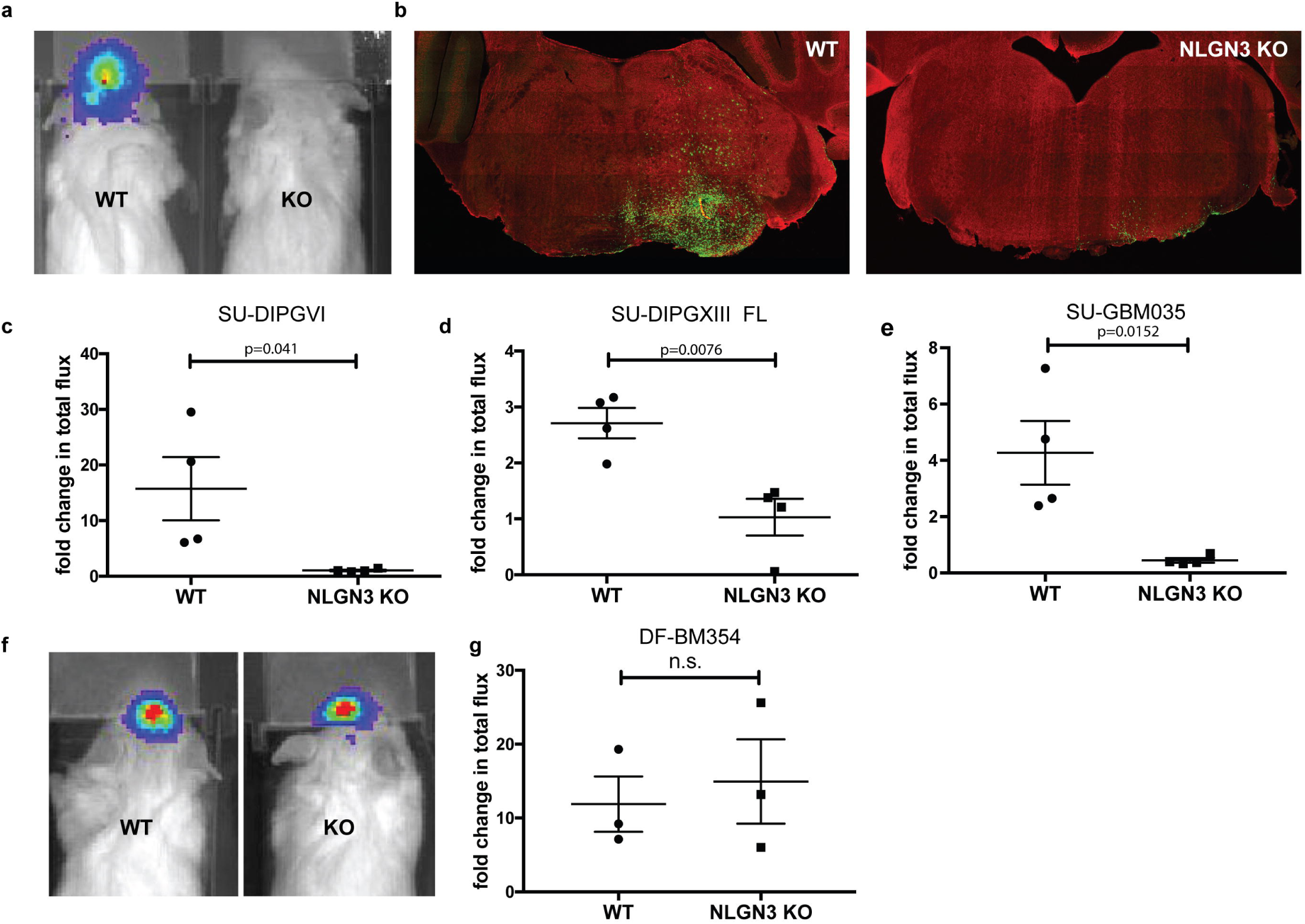
Microenvironmental Nlgn3 is necessary for DIPG and adult GBM growth, but not a breast cancer brain metastasis. Patient-derived DIPG cells (SU-DIPG-VI, from a pontine DIPG tumor or SU-DIPG-XIII FL, from a frontal lobe DIPG metastasis), adult glioblastoma cells, (SU-GBM035 cells) or HER2+ breast cancer brain metastasis cells (DF-BM354 cells) expressing firefly luciferase were xenografted into the orthotopic location of derivation in *Nlgn3*^y/+^;NSG and *Nlgn3*^y/-;^NSG mice. **a,** Representative IVIS images of WT (left) and *Nlgn3* KO (right) mice at 6 weeks following DIPG (SU-DIPG-VI) xenografting. **b,** Representative confocal images at the level of the pons in *Nlgn3* WT (left) and *Nlgn3* KO (right) mouse brains (MBP, red) bearing DIPG xenografts (green) at 6 weeks post-xenografting. **c-e, g**, Fold change in photon emission in SU-DIPG-VI (**c**), SU-DIPG-XIII FL (d), SU-GBM035 (e) and DF-BM354 (g) xenografts in WT and *Nlgn3* knockout (KO) mice at 6 weeks (c,d) or 4 weeks (e,g) after xenografting. **f**, Representative IVIS images of WT (left) and *Nlgn3* KO (right) mice at 4 weeks following breast cancer brain metastasis (DF-BM354) xenografting. *n*=3-4 mice per group. Each dot represents one mouse. P values indicated on graphs, Student’s two-tailed t-test. Data shown as mean+/-s.e.m.

## Neuroligin-3 signaling in glioma cells

The stagnation of growth observed in the Nlgn3-deficient brain is more robust than can be explained by the known effects of Nlgn3 on glioma cell PI3K-mTOR signaling^1^ alone. We therefore sought to better delineate the signaling consequences of Nlgn3 exposure in glioma. First, we utilized phophoproteomic analyses (phosphoantibody array) of pediatric glioblastoma (SU-pcGBM2) cells at 5 and 30 minutes following the addition of recombinant NLGN3. Phosphorylation events upregulated by ~1.3 fold or more following NLGN3 exposure are summarized in Figure 3a-b. These analyses demonstrated early phosphorylation of focal adhesion kinase (FAK) and numerous phosphorylation events classically downstream of FAK, including activation of the SRC kinase cascade, activation of the PI3K-mTOR cascade, and activation of the SHC-RAS-RAF-MEK-ERK cascade (Fig. 3a, b). Additional oncogenic proteins exhibiting increased phosphorylation following NLGN3 exposure include integrin β3, growth factor receptors EGFR, FGFR and VEGFR, as well as others (Fig. 3a, b). Central to many of these signaling consequences of NLGN3 exposure is FAK activity. Phospho-tyrosine pull-down analysis at 5 minute following NLGN3 exposure similarly demonstrated FAK phosphorylation. FAK inhibition blocks the effects of NLGN3 exposure on glioma cell proliferation (Fig. 3c, d). A time course analysis of FAK phosphorylation indicated peak phosphorylation at 5-10 minutes following NLGN3 exposure, indicating this is an early signaling event (Fig. 3e). We had previously shown that PI3K-mTOR signaling is necessary for NLGN3-induced glioma proliferation^1^ and FAK can be upstream of PI3K pathway. Accordingly, we found that FAK activity is necessary for PI3K stimulation by NLGN3 as FAK inhibition abrogates NLGN3-induced phosphorylation of AKT (Fig. 3f).

**Figure 3.**
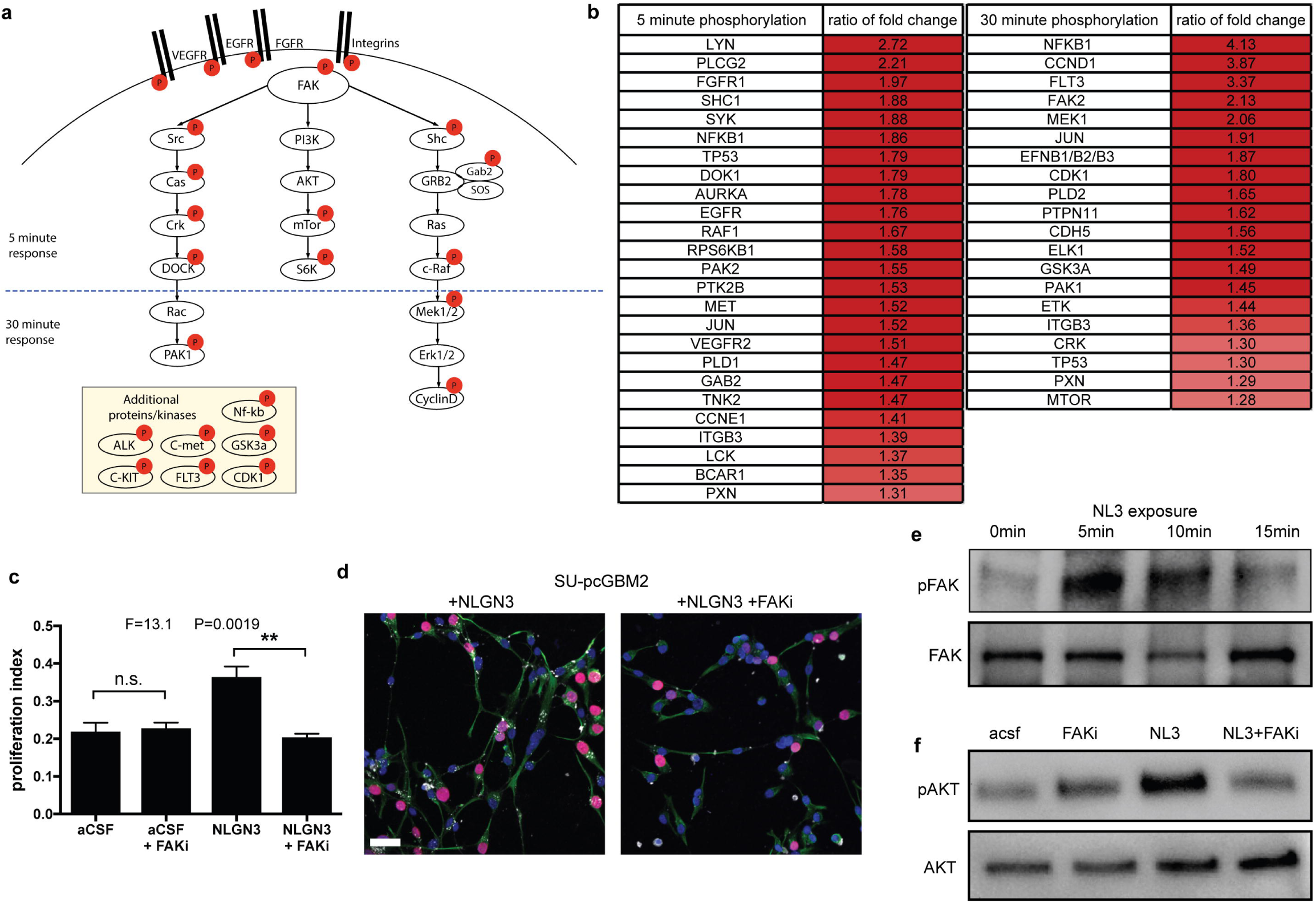
Signaling consequences of NLGN3 in glioma. **a,** Schematic illustration of signaling pathways activated following NLGN3 exposure; red circles represent proteins exhibiting increased phosphorylation following NLGN3 exposure. **b**, Phosphorylation ratio of NLGN3-exposed vs. control SU-pcGBM2 cell lysates after 5-minute or 30-minute exposure to NLGN3 as assessed by phospho-antibody array. **c**, Proliferation index of SU-pcGBM2 cells exposed to plain media (aCSF), aCSF + 5nM FAK inhibitor (FAKi), NLGN3 (50nM), or NLGN3 (50nM) + 5nM FAK inhibitor (NLGN3 + FAKi), (n=3 wells per condition; P and F values indicated on graph; One-way ANOVA, Data shown as mean+/-s.e.m.). **d,** Representative confocal images of SU-pcGBM2 cells exposed to NLGN3 in the absence (left) or presence of FAK inhibitor (right). Vimentin, green; phospho-FAK, white; DAPI, blue; EdU, red, scale bar = 50 μm. **e,** Representative Western blot demonstrating increased phosphorylation of FAK (Tyr861) after 0, 5 or 15-minute exposure to NLGN3. **f**, Representative Western blots demonstrating AKT phosphorylation in SU-pcGBM2 cells exposed to plain media (aCSF), aCSF + 5nM FAK inhibitor (FAKi), NLGN3 (50nM), or soluble NLGN3 (50nM) + 5nM FAK inhibitor, (NLGN3 + FAKi).

To further understand the mechanisms of NLGN3-induced glioma growth, we next performed RNA sequencing (RNA-seq) of pediatric cortical glioblastoma (SU-pcGBM2) cells 16 hours following NLGN3 exposure (Extended Data Fig. 5). As expected, a host of genes involved in cell proliferation were upregulated (Extended Data Fig. 5). We also found significant upregulation of genes known to promote malignancy in glioma, including increased expression of platelet-derived growth factor alpha (PDGFA), a growth factor strongly implicated in glioma growth and progression^13–15^, and several potassium channel genes, recently shown to play a role in DIPG cell viability^16^. In addition to the expected upregulation of NLGN3 expression itself^1^, a number of genes involved in synapse function were also upregulated following NLGN3 exposure (Extended Data Fig. 5). While the meaning of this intriguing observation is not yet clear, it suggests that the biology of NLNG3 in glioma may be more complex than simply promoting proliferation.

## Cellular sources of secreted neuroligin-3

The dramatic glioma growth inhibition observed in the Nlgn3-deficient brain highlights its potential as a therapeutic target. One therapeutic strategy would be to block its release into the tumor microenvironment, and therefore we sought to determine how, and from which cell types, Nlgn3 is cleaved and secreted. As previously shown and re-demonstrated here, full length Nlgn3 is cleaved and secreted in an activity-regulated fashion by enzymatic cleavage and shedding of the N-terminal ectodomain^1^ (Fig. 4a, b). Optogenetic stimulation of acute cortical slices from *Thy1*::ChR2 mice illustrates increased Nlgn3 shedding detected in the CM using western blot analysis as compared to the CM generated from slices exhibiting baseline spontaneous neuronal activity. Conversely, the addition of tetrodotoxin (TTX), a specific voltage-gated sodium channel blocker that abolishes neuronal action potentials, inhibits Nlgn3 secretion in the CM (Fig. 4c).

**Figure 4:**
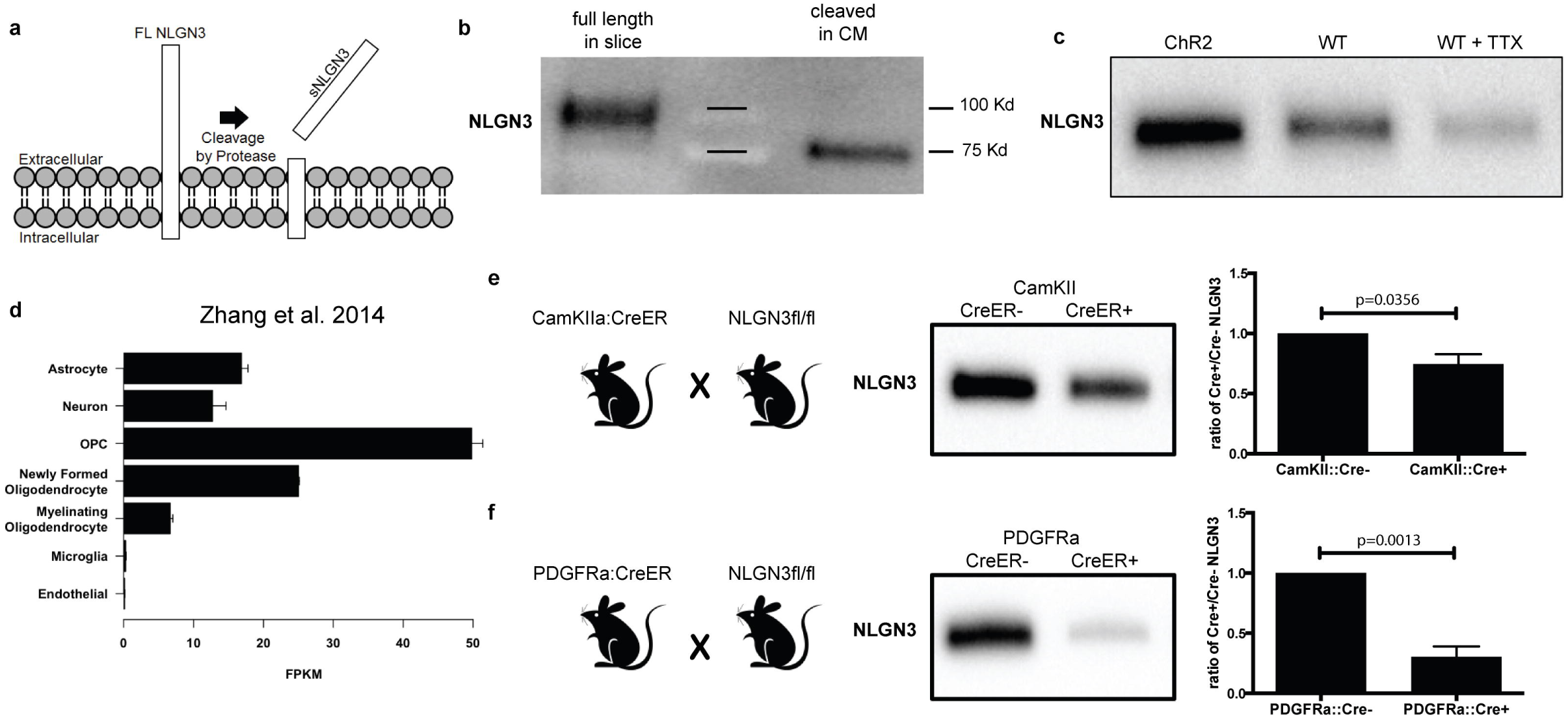
Activity-regulated neuroligin-3 secretion from both neurons and OPCs. **a,** Schematic illustration of proteolytic cleavage of neuroligin-3 at the membrane. **b**, Representative Western blot showing full length Nlgn3 in slice lysate and cleaved Nlgn3 N-terminal ectodomain fragment in conditioned media (CM). **c,** Representative Western blot illustrating secreted Nlgn3 levels in CM from *Thy1*:*:ChR2* cortical slices that have been optogenetically stimulated, WT slices at baseline neuronal activity, and WT slices in the presence of 1 μM tetrodotoxin (TTX). **d,** *Nlgn3* RNA expression (FPKM values) in various cell types; (data from Brain-seq Barres dataset^4^) **e,** *CamKIIa*::Cre^ER^;*Nlgn3*^fl/fl^ mouse model; Western blot analysis and quantification of CM generated from *CamKIIa*::Cre^ER^;*Nlgn3*^fl/fl^ or *Nlgn*3^fl/fl^ (no Cre) cortical slices at baseline neuronal activity. **f,** *PDGFRa*::Cre^ER^;*Nlgn3*^fl/fl^ mouse model; Western blot analysis and quantification of CM from *PDGFRa*::Cre^ER^;*Nlgn3*^fl/fl^ or *Nlgn3*^fl/fl^ (no Cre) cortical slices at baseline neuronal activity. Note that the Western blots in panels e and f were run on the same gel. P values indicated on graphs; Student’s two-tailed t-test. Data are shown as mean+/-s.e.m.

Nlgn3 is expressed at high levels in both neurons and in oligodendrocyte precursor cells (OPCs; Fig. 4d)^4^, known to serve as a post-synaptic cell and form bona fide glutamatergic and GABAergic synapses with presynaptic neurons^17–19^. To test the relative contribution of neurons and OPCs to Nlgn3 secretion, mice expressing either the neuron-specific inducible Cre driver *CamKIIa*::Cre^ER^ or the OPC-specific inducible Cre driver *PDGFRa*::Cre^ER^ were bred to Nlgn3^fl/fl^ mice. Tamoxifen was administered for 5 days beginning at P28, resulting in recombination in ~40% of cortical neurons in the *CamKIIa*::Cre^ER^ driver mouse and in ~80% of OPCs in the *PDGFRa*::Cre^ER^ driver mouse (Extended Data Fig. 6). Using the acute slice paradigm together with inducible, conditional deletion of Nlgn3 from either neurons or from OPCs, we find that both cells types contribute to activity-regulated Nlgn3 secretion, and notably OPCs are a major source of secreted Nlgn3 (Fig.4e, f).

We previously demonstrated that NLGN3 exposure results in feed-forward glioma cell expression of NLGN3 mRNA and protein^1^, and thus asked what contribution glioma cells make to the pool of secreted NLGN3 in the microenvironment. We find elevated levels of secreted neuroligin-3 in CM from glioma xenograft-bearing *Nlgn3* WT brain slices compared to non-xenograft bearing slices (Extended Data Fig S7a-b), either due to NLGN3 secreted from the glioma cells or because glioma cells can increase cortical excitability^20^. To confirm that glioma cells are themselves secreting NLGN3, we collected CM from glioma cells that had been exposed to recombinant NLGN3 or to control vehicle and then washed prior to CM collection. We find up-regulation of NLGN3 secretion from glioma cells following NLGN3 exposure (Extended Data Fig. 7c). Consistent with this finding and underscoring the importance of microenvironmental NLGN3 to stimulate feed-forward glioma cell expression and secretion, xenograft-bearing brain slices from *Nlgn3* KO mice secrete no detectable protein (Extended Data Fig. S7b). Taken together, these data indicate that glioma cells contribute to the secreted NLGN3 in the tumor microenvironment in a manner regulated by neuroligin-3 exposure from normal stromal cells.

## ADAM10 cleaves Neuroligin-3

To determine the enzyme responsible for the cleavage of Nlgn3, the amino acid sequence of the c-terminal region of Nlgn3 was analyzed for putative cleavage sites (https://prosper.erc.monash.edu.au/home.html). The analysis yielded both the MMP and ADAM family proteases as candidate enzymes involved in Nlgn3 secretion, a prediction that is consistent with the reported enzyme(s) responsible for Nlgn1 cleavage^5,6^ and a database of neural ADAM10 targets^21^. To begin to determine the identity of the enzyme responsible for activity-dependent Nlgn3 cleavage, optogenetically stimulated *Thy1*::ChR2 cortical slices were utilized to generate conditioned media in the absence and presence of various protease inhibitors. Western blot analyses of the resulting CM showed a decrease in the level of Nlgn3 in CM with a pan-MMP inhibitor, an MMP2/9 inhibitor, an MMP9/13 inhibitor, and a specific ADAM10 inhibitor^22^ (Fig. 5a). An increase in full-length Nlgn3 was observed in brain slice lysate concomitant with the observed decrease in cleaved Nlgn3 in CM when either MMP9 or ADAM10 inhibitors were used (Fig. 5b). While these results suggest a possible role for MMP9 and/or ADAM10, one MMP9-specific inhibitor had no effect (Fig. 5a). To test the functional effect of these protease inhibitors, HGG cells were exposed to CM generated in the presence of each compound. As expected, the increased proliferation seen in response to active CM was abrogated in the cells exposed to CM generated in the presence of the protease inhibitors that reduce Nlgn3 secretion (Fig. 5c-d). Neither MMP nor ADAM10 inhibitors exerted direct effects on glioma cell proliferation (Fig. 5c-d). Thus, decreased Nlgn3 levels in the CM results in a corresponding decrease in the proliferation index of HGG cells exposed to this CM. These data highlight MMP9 and ADAM10 as promising candidate enzymes mediating activity-dependent NLGN3 cleavage and secretion.

**Figure 5:**
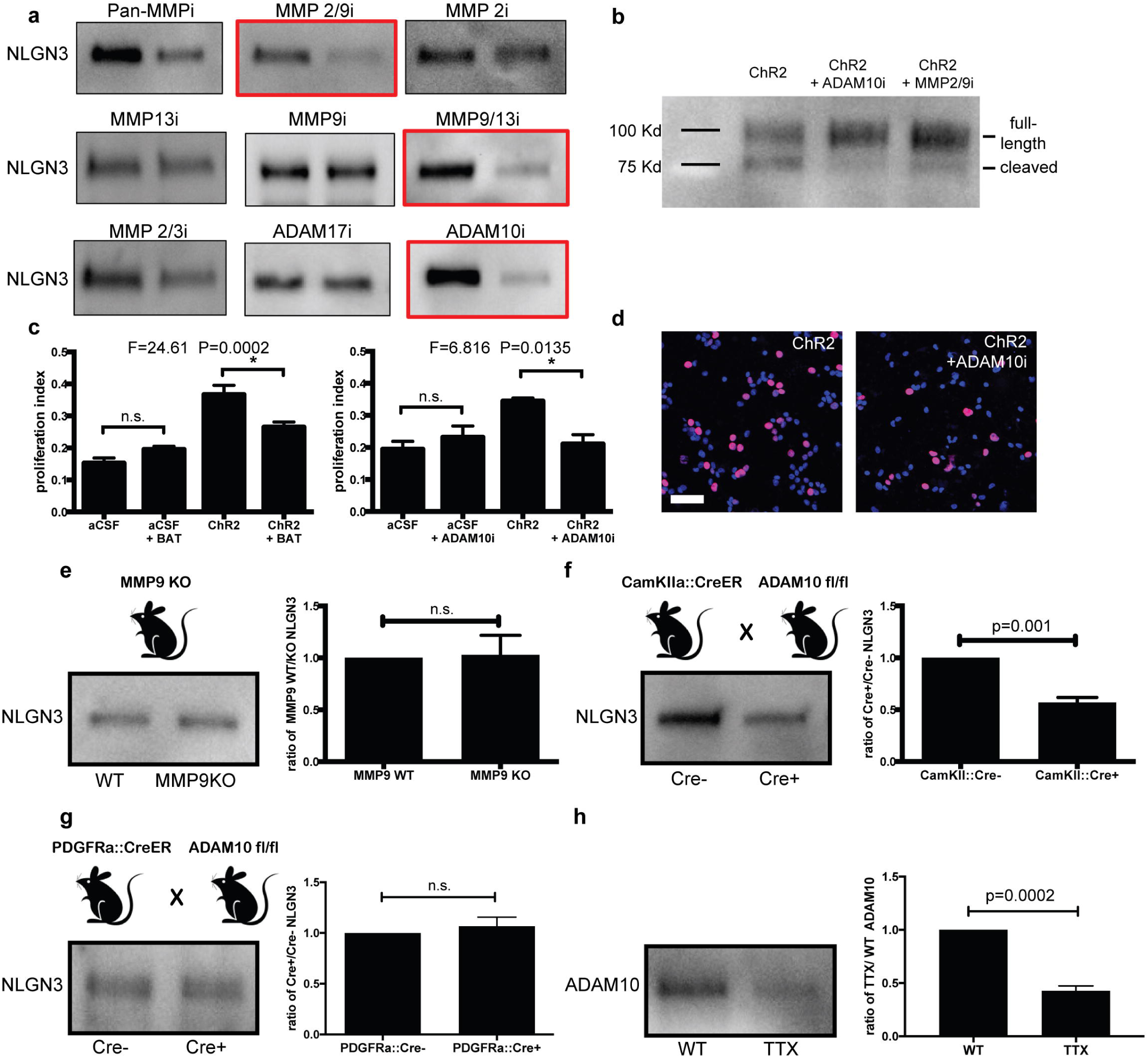
ADAM10 is the primary sheddase involved in neuroligin-3 cleavage. **a,** Representative Nlgn3 Western blot analyses of CM generated from optogenetically stimulated *Thy1*::ChR2 slices in the presence and absence of specific protease inhibitors as indicated. **b,** Nlgn3 Western blot analyses of optogenetically stimulated *Thy1:: ChR2* cortical slice homogenates +/- ADAM10 and MMP2/9 inhibitors. **c,** Proliferation indices of SU-pcGBM2 cells exposed to plain media (aCSF) or CM in the present of absence of pan-MMP inhibitor (BAT, left) or ADAM10 inhibitor (right). n=3 wells per condition. **d,** Representative confocal micrographs of SU-pcGBM2 cells (EdU, red; DAPI, blue) exposed to CM +/- ADAM10 inhibitor; Scale bar = 50μm. **e,** *MMP9*^-/-^ mouse model; Nlgn3 Western blot analysis and quantification illustrating secreted Nlgn3 levels from WT and *MMP9*^-/-^ cortical slices CM at baseline neuronal activity, expressed as ratio of Nlgn3 levels in *MMP9* KO to WT slice CM. **f,** *CamKIIa*::Cre^ER^;*ADAM10*^fl/fl^ mouse model; Nlgn3 Western blot analysis and quantification of CM from *CamKIIa*::Cre^ER^;*ADAM10*^fl/fl^ and *ADAM10^fl/fl^* (no cre) cortical slices at baseline neuronal activity expressed as ratio of Nlgn3 levels in Cre+ to Cre**-** slice CM. **g,** as in f but with *PDGFRa*::Cre driver mouse. **h,** ADAM10 Western blot analysis and quantification of cortical slice CM in the present or absence of TTX. n= 3 biological replicates (e-h). P values shown in graphs; one-way ANOVA (c); two-tailed Student’s t-test (e-h). Data shown as mean+/-s.e.m.

Given the possibility of cross-inhibition with pharmacological protease inhibition, genetic knockout models were employed to more definitively determine whether MMP9 or ADAM10 is responsible for Nlgn3 cleavage and shedding. Acute brain slices from *Mmp9* knockout mice showed no change in Nlgn3 secretion into CM as compared to those from *Mmp9* WT mice (Fig. 5e), suggesting that MMP9 is not responsible for Nlgn3 cleavage and that the decreased levels of Nlgn3 secretion observed with pharmacologic MMP9 inhibition may be due to cross-inhibition/off-target effects. Constitutive knockout of *Adam10* is embryonic lethal, thus an inducible conditional *Adam10* knockout model was used to delete *Adam10* from *CamKIIa-*expressing neurons using the *CamKIIa*::CreER driver mouse as above. Western blot analysis of the CM generated from *CamKIIa*::CreER+; *Adam10*^fl/fl^ cortical slices demonstrated ~50% decrease in secreted Nlgn3 in comparison to *CamKIIa*::CreER-; *Adam10*^fl/fl^ mice (Fig. 5f). In contrast, conditional deletion of *Adam10* from OPCs using *PDGFRA*::CreER; *Adam10*^fl/fl^ mice does not influence Nlgn3 secretion (Fig. 5g). ADAM10 can be released from neurons at synapses where it can act on the ectodomains of synaptic proteins^23^. We find that brain slice incubation with TTX to silence spontaneous neuronal activity substantially abrogates ADAM10 release into slice CM (Fig. 5h), demonstrating activity-regulated release of the ADAM10 sheddase. Taken together these data indicate that ADAM10, released in activity-dependent fashion from neurons, is the chief enzyme responsible for cleaving Nlgn3, and may be a promising target for HGG therapy.

## ADAM10 inhibition blocks glioma growth *in vivo*

As ADAM10 is a key mediator of Nlgn3 secretion, and we have shown the importance of Nlgn3 for high-grade glioma growth *in vivo,* we investigated the therapeutic potential of ADAM10 inhibition for HGG. ADAM10 is expressed in gliomas as reported in the literature^24,25^ and demonstrated in gene expression datasets from pediatric and adult HGG samples^1,16,26–28^ (Extended Data Fig. 7d). Cell intrinsic effects of ADAM10 inhibition have been reported in adult HGG ^25,29,30^, so prior to assessing possible effects of modulating Nlgn3 secretion in the glioma microenvironment with ADAM10 inhibition we first sought to assess possible direct effects of this inhibitor on the pHGG cells used here. Cell-intrinsic effects of ADAM10 inhibition were not observed on proliferation, overall cell viability or invasion of pHGG cells, even at high concentrations (Fig. 6a, b Extended Data Fig. 8a). We did find a mild effect of ADAM10 inhibition on pHGG self-renewal in a neurosphere formation assay (Fig. 6c and Extended Data Fig. 8b), consistent with reports in adult HGG^25^.

**Figure 6:**
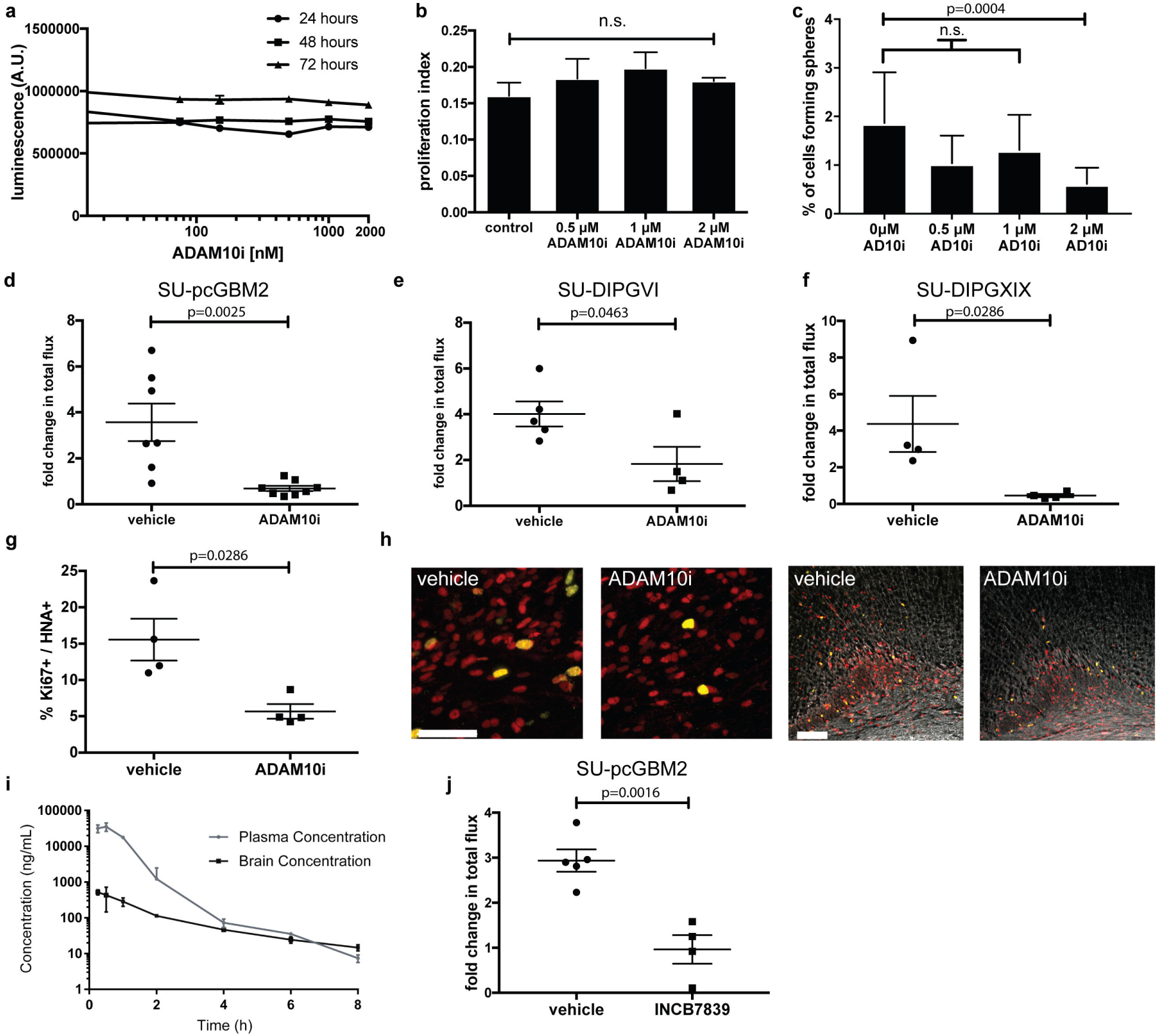
ADAM10 inhibition blocks glioma growth. **a,** Cell viability of SU-pcGBM2 cells exposed to ADAM10 inhibitor (GI254023X, 10nM-2μM) or vehicle control at 24-, 48-, and 72-hours, (n=3 wells/condition) **b**, Proliferation index of SU-pcGBM2 cells exposed to GI254023X (0-2μM; (n=3 wells/condition) **c**, Neurosphere formation assay in SU-pcGBM2 cells in the presence of GI254023X (0-2μM; (n=10 wells/condition)) **d-f,** Growth (change in photon emission) of orthotopic xenografts following systemic administration of GI254023X or vehicle control for SU-pcGBM2 (d), SU-DIPG-VI (e), SU-DIPG-XIX (f) xenografts (*n* = 4-7 mice/group.) **g**, *In vivo* proliferation index of SU-pcGBM2 cells in vehicle control and ADAM10i treated mice (*n*=4/group). **h,** Representative confocal images (Ki67^+^, green; human nuclear antigen, HNA^+^ red) of vehicle-treated or ADAM10i-treated mice. Scale bars=50 μm. **i,** Brain and tissue penetration of INCB7839; (n=3 mice/data point). **j,** Growth (change in photon emission) of SU-pcGBM2 xenografts following systemic administration of INCB7839 or vehicle control; (*n* = 4-5 mice/group.) P values indicated in graphs. n.s. = not significant. one-way ANOVA (a,b), *χ*^2^ test (c), Student’s two-tailed t-test (d,e,j,); Mann-Whitney (f,g). Each dot represents one mouse. Data are shown as mean +/- s.e.m except in c) error bars are +/- confidence intervals and in i) error bars are st. dev.

We next tested the influence of ADAM10 inhibition on HGG growth *in vivo*. Brain penetration of the specific ADAM10 inhibitor GI254023X was assessed following systemic administration using liquid chromatography/tandem mass spectrometry (LC-MS/MS). Following a single 100 mg/kg intraperitoneal (i.p.) dose, drug levels in brain tissue were found to be ~2-4 μM, indicating sufficient blood-brain-barrier penetration (Extended Data Table 1). Given the possible effects of ADAM10 inhibition on glioma self-renewal and therefore tumor initiation, drug treatment started well after engraftment had occurred. Pediatric glioblastoma (SU-pcGBM2-GFP-luc) cells were xenografted into the cortex or DIPG (SU-DIPGVI-GFP-luc or SU-DIPG-XIX) cells were xenografted into the pons of NSG mice. Four weeks after xenografting, mice were treated with either the ADAM10 inhibitor GI254023X (100 mg/kg) or vehicle control 5 times/week over the course of 2 weeks and monitored using *in vivo* bioluminescent imaging. GI254023X was well tolerated, with no apparent adverse events. The pGBM and both DIPG xenografts in mice treated with the ADAM10 inhibitor exhibited a robust reduction in growth as compared to vehicle-treated controls (Fig. 6d-f), commensurate with the stagnation of glioma xenografts in the Nlgn3-deficient brain described above. Histological analyses revealed a reduced proliferation index in ADAM10 inhibitor-treated animals as assessed by the fraction of glioma cells co-expressing the proliferation marker Ki67 (Fig. 6g-h). Furthermore, we found that ADAM10 inhibition abrogates glioma cell secretion of neuroligin-3 from both xenograft-bearing brain slices and NLGN3-primed glioma cells *in vitro* (Extended Data Fig. 7b-c), suggesting this therapeutic strategy addresses all cellular sources of neuroligin3.

ADAM10 inhibitors have been developed for clinical use, and the ADAM10/17 inhibitors INCB7839 (aderbasib) and XL-784 have been through Phase II clinical trials for indications such as breast cancer^31,32^. We used LC-MS/MS to evaluate brain penetration of these compounds, and found the pharmacokinetic profile of INCB7839 was superior to XL-784 (Fig. 6i and Extended Data Fig. 9). INCB7839 inhibits ADAM10 enzymatic function at low nanomolar concentrations^31,32^ and was found to reach a brain tissue level of 100 ng/ml (~240 nM) for ~4 hours following systemic administration in mice, suggesting sufficient BBB penetration (Fig. 6i). We therefore tested INCB7839 at a clinically relevant dose (50 mg/kg) administered 5 days per week beginning four weeks following xenografting and found that, like GI254023X, INCB7839 robustly inhibits growth of pediatric glioblastoma orthotopic xenografts (Fig. 6j). Targeting of the sheddase ADAM10 thus represents a strategy to modulate neuroligin-3 levels in the tumor microenvironment for HGG therapy.

## Discussion

Here, we demonstrate an unexpected and conserved dependency of HGG growth on microenvironmental neuroligin-3 across multiple classes of pediatric and adult HGG. The magnitude of this effect both underscores its potential as a therapeutic target and suggests that the mechanisms by which NLGN3 promotes HGG growth are incompletely understood. We have furthered our understanding of the mechanisms by which NLGN3 stimulates glioma proliferation and now show that NLGN3 activates FAK upstream of the PI3K-mTOR pathway, as well as stimulating numerous additional oncogenic signaling pathways and upregulating genes associated with known mechanisms of glioma malignancy. However, loss of a single robust mitogen is unlikely to account entirely for the complete stagnation of xenograft growth observed in the Nlgn3-deficient brain. Future work will need to elucidate further mechanisms by which NLGN3 regulates glioma progression as well as by which the cancer may circumvent this dependency.

NLGN3 is cleaved and secreted from both neurons and OPCs. The physiological function of NLGN3 cleavage remains to be determined. Whether activity-regulated cleavage of NLGN3 in OPCs, likely occurring at axo-glial synapses, plays a role in the adaptive responses of myelin-forming precursor cells to neuronal activity^34^ remains to be seen. However, these findings define a previously unrecognized place for OPCs as a microenvironmental cell type contributing to glioma growth.

Targeting NLGN3 holds great promise for glioma therapy as an adjuvant to traditional cytocidal treatment modalities such as radiation and chemotherapy. Possible strategies to prevent the glioma growth-promoting effects of NLGN3 include blocking NLGN3 cleavage and release, sequestering soluble NLGN3 in the tumor microenvironment or blocking the as-of-yet unidentified NLGN3 binding partner/receptor on glioma cells. Here, we focus on the former strategy, identify ADAM10 as the protease mediating NLGN3 cleavage and demonstrate robust stagnation of tumor growth with ADAM10 inhibition in preclinical models of HGG. The specific ADAM10 inhibitor GI254023X is presently in the preclinical phase of development, but the ADAM10 inhibitor INCB7839 has been through phase II clinical testing for other indications and could be re-purposed now for HGG therapy. ADAM10 mediates cleavage of numerous cell surface proteins, prominently targeting synapse-associated proteins^21^, and also plays an important role in amyloid protein processing^33^. While ADAM10 inhibition appears well-tolerated in clinical trials^31,32^, long-term effects on synaptic plasticity, amyloid deposition and neurological function should be carefully evaluated as part of any effort towards clinical translation of ADAM10 inhibition for this group of lethal cancers.

The present study highlights a means to target the growth-promoting effects of neuronal activity on high-grade gliomas that span a range of clinically and molecularly distinct glioma types. Further elucidation of neural influences on other classes of brain cancers may provide additional therapeutic insights. Moreover, a critical role is emerging for nervous system regulation of a variety of cancers, including prostate^35^, gastric^36^, pancreatic^37^ and skin^38^ cancers. Deeper understanding of the neural mechanisms driving malignancy may broadly highlight avenues towards more effective cancer control.

## Methods

### Mice and housing conditions

All *in vivo* experiments were approved by the Stanford University Institutional Animal Care and Use Committee and performed in accordance with institutional guidelines. Animals were housed according to standard guidelines with free access to food and water in a 12-hour light/dark cycle.

For constitutive knockout studies, *Nlgn3* KO mice (The Jackson Laboratory) were intercrossed with NSG mice (NOD-SCID-IL2R gamma chain-deficient, The Jackson Laboratory) to produce the *Nlgn3* KO;NSG genotype. All experiments were performed on male mice either hemizygous WT *Nlgn3* or hemizygous null *Nlgn3* littermates.

For conditional knockout experiments, *Nlgn3*^fl/fl^ mice (a kind gift from Dr. Thomas Sudhof) or *ADAM10*^fl/fl^ mice (The Jackson Laboratory) were crossed to *CamKIIa*:CreER (The Jackson Laboratory) or *PDGFRa*::CreER (The Jackson Laboratory). Cre+ or control Cre- floxed mice were treated with 100mg/kg tamoxifen i.p. for 5 days and experiments were performed 7 days after the end of treatment. Tamoxifen was given from postnatal day 28 (P28) to P33 and slice experiments performed at P40.

### Bioluminescent imaging

For *in vivo* monitoring of tumor growth, bioluminescence imaging was performed using an IVIS imaging system (Xenogen). Mice orthotopically xenografted with luciferase-expressing glioma cells were placed under isofluorane anesthesia and injected with luciferin substrate. Animals were imaged at baseline and randomized based on tumor size by a blinded investigator so that experimental groups contained an equivalent range of tumor sizes. All total flux values were then normalized to baseline values to determine fold change of tumor growth. Statistical analysis between tumor burden in each group was assessed using Student’s two-tailed t-test (parametric data) or Mann Whitney test (non-parametric data). Based on the variance of xenograft growth in control mice, we used at least 3 mice per genotype to give 80% power to detect an effect size of 20% with at a significance level of 0.05.

### Orthotopic xenografting

A single-cell suspension from cultured neurospheres was prepared in sterile PBS immediately prior to the xenograft procedure. Animals at p34-36 were anesthetized with 1-4% isoflurane and placed in a stereotactic apparatus. The cranium was exposed via midline incision under aseptic conditions. 600,000 glioma cells in 3 μL sterile PBS were stereotactically implanted in either the premotor cortex (M2) of the right hemisphere (SU-pcGBM2) or the pons (SU-DIPGVI and SU-DIPGXIX) through a 31-gauge burr hole, using a digital pump at infusion rate of 0.4 μL/minute and 31-gauge Hamilton syringe. For breast cancer brain metastasis studies, 100,000 DF-BM354 cells were similarly xenografted in the premotor cortex. Stereotactic coordinates used were as follows: for premotor cortex, 0.5 mm lateral to midline, 1.0 mm anterior to bregma, –1.75 mm deep to cranial surface; for pons, 1.0 mm lateral to midline, -0.8 mm posterior to lamda, –5.0 mm deep to cranial surface. At the completion of infusion, syringe needle was allowed to remain in place for a minimum of 2 minutes, then manually withdrawn at a rate of 0.875 mm/minute to minimize backflow of the injected cell suspension.

### Perfusion and immunohistochemistry

Animals were anesthetized with intraperitoneal Avertin (tribromoethanol), then transcardially perfused with 20 mL of PBS. Brains were fixed in 4% paraformaldehyde overnight at 4°C, then transferred to 30% sucrose for cryoprotection. Brains were embedded in Tissue-Tek O.C.T. (Sakura) and sectioned in the coronal plane at 40 μm using a sliding microtome (Microm HM450; Thermo Scientific). For immunohistochemistry, coronal sections were incubated in blocking solution (3% normal donkey serum, 0.3% Triton X-100 in TBS) at room temperature for 30 minutes. Chicken anti-GFP (1:500, Abcam), Rat anti-MBP (1:300, Abcam), Mouse anti-human nuclei clone 235-1 (1:100; Millipore), or rabbit anti-Ki67 (1:500; Abcam), were diluted in 1% blocking solution (1% normal donkey serum in 0.3% Triton X-100 in TBS) and incubated overnight at 4°C. Sections were then rinsed three times in 1X TBS and incubated in secondary antibody solution (Alexa 488 donkey anti-chicken IgG, 1:500 (Jackson Immuno Research); Alexa 594 donkey anti-mouse IgG, 1:500 (Life Technologies); Alexa 647 donkey anti-rabbit IgG, 1:500 (Life Technologies); Alexa 594 donkey anti-rat IgG, 1:1000 (Life Technologies)) in 1% blocking solution at 4°C overnight. Sections were rinsed 3 times in TBS and mounted with ProLong Gold Mounting medium (Life Technologies).

### Cell culture

For all human tissue studies, informed consent was obtained and Institutional Review Board (IRB) approval was granted. For all cultures, short tandem repeat (STR) DNA fingerprinting was performed every three months to verify authenticity. The STR fingerprints and clinical characteristics for the patient-derived cultures used have been previously reported^1^ with the exception of SU-DIPG-XIX which is a H3.3K27M mutant tumor that was derived at the time of autopsy from a male who was 2 years of age at diagnosis, was treated with radiotherapy and cabazitaxel and who survived 18 months. STR fingerprint for SU-DIPG-XIX is: X/Y (AMEL), 10/11 (CSF1PO1), 13/14 (D13S317), 9/13 (D16S539), 30/30 (D21S11), 11/12 (D5S818), 10/10 (D7S820), 9.3/9.3 (TH01), 8/11 (TPOX), 17/18 (vWA). All cell cultures were routinely tested for mycoplasma.

All high-grade glioma cultures were generated as previously described^1^. In brief, tissue was obtained from high-grade glioma (WHO grade III or IV) tumors at the time of biopsy or from early post-mortem donations. Tissue was dissociated both mechanically and enzymatically and grown in a defined, serum- free medium designated “Tumor Stem Media (TSM)”, consisting of Neurobasal(-A) (Invitrogen), B27(-A) (Invitrogen), human- bFGF (20 ng/mL) (Shenandoah Biotech), human-EGF (20 ng/mL) (Shenandoah), human PDGF-AA (10 ng/mL) and PDGF-BB (10 ng/mL) (Shenandoah) and heparin (2 ng/mL) (Stem Cell Technologies).

Breast cancer brain metastasis line, PDX DF-BM354, was provided by the Zhao lab and developed as previously described^12^.

### Generation of conditioned media from acute cortical slices

Conditioned media was generated as previously described^1^. Mice (genotype varied based on experiment) between the ages of 4-7 weeks were briefly exposed to 4% isofluorane and immediately cervically dislocated and decapitated. Extracted brains were placed in oxygenated high-sucrose solution and sliced in 350-μm sections. Slices were then placed in buffering solution (aCSF) and allowed to recover for at least one hour. After recovery, slices were then moved into fresh aCSF in a 24-well plate and slices optogenetically stimulated using a blue-light LED to observe the effects of elevated neuronal activity (in the case of *Thy1*::ChR2 brain slices) or unstimulated to observe the effects of baseline neuronal activity. Following recovery, media was conditioned for 30 minutes. For various experiments, conditioned media was prepared in the presence of various protease inhibitors (described below) or tetrodotoxin at 1uM (Tocris). Surrounding medium was then collected for immediate use or frozen at -80°C for future experiments. All slice experiments were performed in three biological replicates unless otherwise indicated.

### EdU incorporation assay

8-well chamber slides were coated with poly-L-lysine. Cells were then seeded at 40,000 cells per well and exposed to aCSF, aCSF the relevant inhibitor (see below), or active conditioned media (conditioning methods vary by assay). 10 μM EdU was added to each well. Cells were fixed after 24 hours using 4% paraformaldehyde in PBS and stained using the Click-iT EdU kit and protocol (Invitrogen). Proliferation index was then determined by quantifying percentage of EdU labeled cells using confocal microscopy at 200x magnification. Group mean differences were otherwise assessed using one-way analysis of variance (one-way ANOVA) with Tukey post-hoc tests to further examine pairwise differences. All experiments were performed in three biological replicates.

### CellTiter-Glo assay of cell viability

To assess overall cell number, 5000 glioma cells were seeded in minimal growth media in a 96-well plate with varying concentrations of ADAM10 inhibitor. After 24, 48, or 72 hours, CellTiter-Glo reagent (Promega) was added at a 1:1 ratio. Luminescence was measured after 10- minute incubation at room temperature to stabilize signal. All experiments were performed in three biological replicates.

### Inhibitors used

Batimastat at 20nM (Pan MMP inhibitor; BB-94; Selleck Chemicals); MMP-2/MMP-9 Inhibitor II at 50nM (sc-311430; Santa Cruz biotechnology); ARP 100 at 20nM (MMP2 inhibitor) (R&D Systems); MMP-13 Inhibitor at 10nM (sc-205756; Santa Cruz biotechnology); MMP-9 Inhibitor I at 100nM (sc-311437; Santa Cruz biotechnology); MMP-9/MMP-13 Inhibitor II at 10nM (sc-311439; Santa Cruz biotechnology); MMP2/MMP-3 Inhibitor I at 20uM (sc-295483; Santa Cruz biotechnology); TAPI-1 at 20uM (ADAM17 inhibitor; S7434; Selleck Chemicals); GI 254023X at 200nM (ADAM10 inhibitor; Sigma Aldrich); PF-00562271 at 5nM (FAK inhibitor; S2672; Selleck Chemicals).

### Analysis of NLGN3 secretion from glioma

For *in vivo* studies demonstrating NLGN3 secretion from xenograft-bearing slices, mice were xenografted as above with SU-DIPGXIII or SUGBM035 cells in premotor cortex. Brains were extracted and used for conditioned media experiments in comparison to non-xenografted littermate controls at 5 months (SU-DIPG-XIII) or 6 weeks (SU-GBM035). These experiments were performed in duplicate for SU-DIPG-XIII xenograft-bearing cortical slices and in triplicate for SU-GBM035 xenograft-bearing cortical slices (five biological replicates in total). For *in vivo* studies demonstrating neuroligin secretion from xenograft-bearing slices can be blocked by ADAM10 inhibition, mice were xenografted as above with SU-GBM035 cells in premotor cortex and brains were extracted at 6 weeks post xenograft. Cortical slices were made and incubated in the presence or absence of 200nM ADAM10 inhibitor, GI 254023X. CM was then analyzed using western blot analyses, comparing non-xenograft bearing slices to xenograft-bearing slices from WT or Nlgn3 KO mice in the presence of absence of ADAM inhibitor. These experiments were performed in triplicate.

For *in vitro* studies of NLGN3 secretion from glioma cells, SU-pcGBM2 cells were seeded at 5 million cells/well in the presence of either vehicle, 100nM recombinant NLGN3, 200nM GI 254023X, or 100nM NLGN3 + 200nM GI 254023X for 48 hours. After thorough rinsing of the cells, cells were left in either fresh media or 200nM GI 254023X for another 48 hours in the presence or absence of ADAM10 inhibition. After 48 hours, media was collected and analyzed for presence of cleaved NLGN3 using western blot analyses as described below. Experiments were performed in three biological replicates.

*Spheroid invasion assay* was performed as previously described^16.^

### Neurosphere formation assay

ELDA (extreme limiting dilution analysis) was performed to evaluate self-renewal capacity^39^. SU-pcGBM2 cells were dissociated in TrypLE (+ DNase and HEPES) for 5min at 37C. Cells were triturated into a single cell solution. The solution was incubated with Hoechst (Thermo, cat. 33342) for 30min at 37C. Live cells were identified using a LIVE/DEAD staining kit (Thermo, cat. L10119). Live cells were sorted into 96 well plates. Spheres were counted at 14 days. Cell density per well ranged from: 1, 10, 25, 50, 100, 250, 500, 1000. Each condition was tested in 10 independent wells. Volume of media per well was 200 ul media with growth factors spike-ins every 3-4 days. The ADAM10 inhibitor was reconstituted in DMSO. Each well was adjusted to have 0.1% DMSO, except for the no-DMSO control wells. Neurosphere-forming capability was determined using the Extreme Limiting Dilution Analysis (ELDA) web-based tool (http://bioinf.wehi.edu.au/software/elda/).

### Western Blot analyses

Western blot analyses were used to probe for protein levels present in either brain slice homogenate or secreted into slice conditioned media. For slice homogenates, brain slices were lysed using RIPA buffer and protease inhibitors. Lysates were incubated on ice for 10 minutes and then centrifuged for 10 minutes at 4°C. All samples were normalized to protein concentration, mixed with Laemmli loading buffer (1:4), boiled for 5 minutes, and loaded onto BioRad Mini-Protean TGX precast gels. Protein was transferred to PVDF membranes and blocked with 5% bovine serum albumin (BSA) in TBST for one hour. Anti-Neuroligin-3 (NovusBio; 1:300), Anti-phospho FAK pTyr861 (Thermo Fisher Scientific; 1:500), and anti-FAK (Cell Signaling Technologies; 1:500), or anti-ADAM10 (Abcam; 1:500) were diluted in 1% BSA/TBST and incubated with the membrane overnight. Secondary anti-rabbit conjugated to HRP (BioRad) was then added for one hour (1:1000). Proteins were visualized using Clarity ECL Western Substrate (BioRad) and quantified and analyzed using ImageJ.

### Phospho-antibody array

The Phospho Explorer antibody array assay was performed by Full Moon Biosystems on patient-derived pediatric glioblastoma (SU-pcGBM2) cells. Cell lysates were prepared using Protein Extraction Kit (Full Moon BioSystems). Clear supernatant of the lysates was separated, biotinylated, and incubated with Phospho Explorer Antibody Arrays (Full Moon BioSystems) for two hours at room temperature. The array slides were washed with Wash Buffer (Full Moon BioSystems) and rinsed with DI water. The slides were then incubated with Cy3-Streptavidin for 45 minutes at room temperature, then washed, rinsed and dried. Arrays were scanned on GenePix Array Scanner (Molecular Devices). Image quantification was performed on GenePix Pro (Molecular Devices). Signal intensity data for each spot on the array was extracted from array images. Since there are two replicated printed for each antibody, the mean signal intensity of the replicates is determined. The data is then normalized to the median value (signal intensity) of all antibodies on the slide. Finally, fold change between Control Sample and Treatment Sample is determined using the normalized data (Treatment Sample’s signal is divided by Control Sample’s signal).

### Phosphotyrosine pull down assay

Samples were analyzed using the Cell Signaling Technology PTMScan method as previously described^40–42^. Cellular extracts were prepared in urea lysis buffer, sonicated, centrifuged, reduced with DTT, and alkylated with iodoacetamide. 15mg total protein for each sample was digested with trypsin and purified over C18 columns for enrichment with the Phosphotyrosine pY-1000 antibody (#8803) and the PTMScan Direct Tyr Kinases Reagent^42^. Enriched peptides were purified over C18 STAGE tips (Rappsilber). Enriched peptides were subjected to secondary digest with trypsin and second STAGE tip prior to LC-MS/MS analysis.

Replicate injections of each sample were run non-sequentially for each enrichment. Peptides were eluted using a 90-minute linear gradient of acetonitrile in 0.125% formic acid delivered at 280 nL/min. Tandem mass spectra were collected in a data-dependent manner with an LTQ Orbitrap Elite mass spectrometer running XCalibur 2.0.7 SP1 using a top-twenty MS/MS method, a dynamic repeat count of one, and a repeat duration of 30 sec. Real time recalibration of mass error was performed using lock mass^43^ with a singly charged polysiloxane ion m/z = 371.101237.

MS/MS spectra were evaluated using SEQUEST and the Core platform from Harvard University^44–46^. Files were searched against the NCBI *rattus norvegicus* FASTA database updated on May 22, 2015. A mass accuracy of +/-5 ppm was used for precursor ions and 1.0 Da for product ions. Enzyme specificity was limited to trypsin, with at least one LysC or tryptic (K- or R-containing) terminus required per peptide and up to four mis-cleavages allowed. Cysteine carboxamidomethylation was specified as a static modification, oxidation of methionine and phosphorylation on serine, threonine, and tyrosine residues were allowed as variable modifications. Reverse decoy databases were included for all searches to estimate false discovery rates, and filtered using a 5% FDR in the Linear Discriminant module of Core. Peptides were also manually filtered using a -/+ 5ppm mass error range and reagent-specific criteria. For each antibody reagent results were filtered to include only phosphopeptides matching the sequence motif(s) targeted by the antibodies included. All quantitative results were generated using Progenesis V4.1 (Waters Corporation) to extract the integrated peak area of the corresponding peptide assignments. Accuracy of quantitative data was ensured by manual review in Progenesis or in the ion chromatogram files.

### RNA Sequencing

Samples were processed and analyzed as previously described^16^ with minor modifications as indicated below:

RNA was extracted from Trizol-lysed cells and 1 μg of total RNA was used for each sample. Polyadenylated RNA was selected using Ambion Dynabeads mRNA Purification Kit (Life Technologies 61006) and fragmented with Fragmentation Buffer (Ambion, #AM8740). First strand synthesis was performed using Random Hexamer Primers (Invitrogen, #48190-011) and SuperScript II (Invitrogen, #18064-014). Second strand synthesis was performed using DNA Pol I (Invitrogen #18010-025) and RNA was removed using RNaseH (Invitrogen #18021-014).

Libraries were end-repaired, 3’ A-tailed, and ligated to NEBNext Multiplex Oligo Adaptors (NEB E7335S). Sequencing was performed on an Illumina NextSeq 500 by Stanford Functional Genomics Facility.

Reads were mapped to hg19 annotation using Tophat2 (PMID: 23618408) (version 2.0.13) and transcript expression was quantified against RefSeq gene annotations using featureCounts^47^. Differential testing and log2 fold change calculation was performed using DESeq2^48^. Gene Ontology analyses were performed using DAVID^49,50^.

### Pharmacokinetic studies

### LC-MS/MS analysis of GI254023X concentrations in tissues and serum

#### Sample preparation

A single 100 mg/kg dose was delivered i.p. in NOD-SCID-IL2R γ chain–deficient mice, and tissue samples collected 30 min later for analysis using liquid chromatography/tandem mass spectrometry (LC-MS/MS). Tissues samples were weighed and 1 volume of bullet blender beads (Next Advance) and 3 volume of Milli-Q water were added. Tissues were homogenized by a bullet blender (Next Advance) at 4°C according to manufacturer’s instruction. The neat stock solution of GI254023X was dissolved in DMSO at 5 mg/ml and further diluted in 50% methanol to prepare spiking solutions. For spiked standard curve, 25 μl of GI254023X spiking solutions (0.5 - 1000 ng/ml for brain samples and 1 – 100 μg/ml for serum and kidney samples) was mixed with 25 μl of blank tissue homogenate or serum. For samples, the spiking solution was replaced by 25 μl of 50% methanol to make up the volume. After vortexing all standards and samples, 150 μl of methanol/acetonitrile 20:80 (v/v) was added to the mixture and vortexed vigorously for 1 min followed by centrifugation at 3,000 g for 10 min. The supernatant was diluted 3 times in Milli-Q water for brain samples and 100 times in 25% methanol for serum and kidney samples. The LC-MS/MS system consists of a QTRAP 4000 mass spectrometer (AB SCIEX) coupled to a Shimadzu UFLC system. LC separation was carried out on a ZORBAX SB-Phenyl column (4.6 mm × 50 mm, 3.5 μm) (Agilent) at room temperature. The analysis time was 3 min. The injection volume was 5-10 μl. Isocratic elution was carried out with a mobile phase composed of 55% water and 45% acetonitrile with 0.1% of formic acid and a flow rate of 0.5 ml/min. The mass spectrometer was operated in the positive mode with multiple-reaction monitoring (MRM) with the transition m/z 392.2→361.2. Data acquisition and analysis were performed using the Analyst 1.6.1 software (AB SCIEX).

#### LC-MS/MS analysis of INCB7839 (Aderbasib) and XL-784 in brain tissue and serum

Aderbasib was purchased from Astatech, Inc. XL-784 was provided by True Pharmachem, Inc. A single 50 mg/kg dose of INCB7839 (aderbasib) or XL-784 was delivered i.p. in NOD-SCID-IL2R γ chain–deficient (NSG) mice, and tissue samples collected at various time points for analysis using liquid chromatography/tandem mass spectrometry (LC-MS/MS). NSG mice were purchased from the Model Animal Research Center of Nanjing University. Study was conducted by Crown Biosciences, Inc. Compounds were formulated in 2% DMSO, 2% Tween 80, 48% PEG300, 48% water. Compounds were administered IP, with a dosing volume of 10 μL/g and concentration of 5 mg/ml. Compounds were dosed at 50 mg/kg. A cohort of 24 male, NSG mice, age 6-8 weeks, body weight 18-22 grams, were used for each study. Animals were housed at room temperature, under 40-70% humidity, with a 12 hour light/12 hour dark schedule. Mice were fed with Co^60^ dry granule food, with free access to reverse osmosis water. Eight time points were collected for each compound (0.25, 0.5, 1, 2, 4, 6, 8, and 24 h), with an n = 3 for each time point. Blood was collected via cardiac puncture and collected into potassium-EDTA Eppendorf tubes. Samples were centrifuged within 30 minutes to afford plasma samples. Brains were collected at each time point, PBS (4X) was added, and the material homogenized with a Tissue Lyser II to give a fine homogenate. Brain homogenate (50 μL) was treated with 250 μL acetonitrile (containing 200 ng/mL tolbutamide), which was then vortexed and then centrifuged at 4000 rpm for 20 min. The supernatant was collected and mixed with a 0.1% aqueous formic acid solution. Samples were analyzed on a Waters UPLC or Agilent 1200 Liquid chromatography system, with an API 4000 mass spectrometer, and a 10 μL injection volume, with tolbutamide as an internal standard. Pharmacokinetics were analyzed using WinNonlin6.3 (non-compartmental model).

#### Mouse drug treatment studies

For all drug studies, NSG mice were xenografted as above with either SU-pcGBM2, SU-DIPGVI, or SU-DIPG-XIX. Four weeks post-xenograft, animals were treated with either systemic administration of ADAM10 inhibitor, GI254023X (Sigma-Aldrich; formulated in 10% DMSO in 0.1M carbonate buffer) via intraperitoneal injection 5 days per week at 100mg/kg or systemic administration of ADAM10/17 inhibitor, Aderbasib (Astatech, Inc; formulated in 2% DMSO, 2% Tween 80, 48% PEG300, 48% water) via intraperitoneal injection 5 days per week at 50mg/kg. For both studies, controls were intraperitoneally injected with an identical volume of vehicle. Drug treatment began four weeks after xenografting and continued through week six. Bioluminescence imaging was performed by a blinded investigator before treatment, and again 7 days and 14 days later, using an IVIS imaging system (Xenogen) under isoflurane anesthesia. Similar to above, tumor burden was assessed as fold change in total flux from the beginning to end of treatment. All differences were statistically verified using Students two-tailed t-test.

#### Confocal imaging and quantification of cell proliferation

Cell quantification was performed by a blinded investigator using live counting at 400x magnification using a Zeiss LSM700 scanning confocal microscope and Zen 2011 imaging software (Carl Zeiss Inc.). The area for quantification was selected as follows: of a 1 in 6 series of 40-μm coronal sections, 3 consecutive sections were selected at approximately 1.1–0.86 mm anterior to bregma (Figures 22, 23, 24; Franklin & Paxinos, *The Mouse Brain in Stereotaxic Coordinates, 3^rd^ Ed.* 2008); using our stereotactic coordinates for tumor xenograft, these sections are expected to include the tissue most proximal to the site of tumor cell implantation in the coronal plane. For each of the three consecutive sections, the cingulum bundle was first identified as an anatomic landmark, and six 160x160-μm field area for quantification were selected centered around this landmark within cortical layer 6b of M2. Within each field, all human nuclear antigen (HNA)-positive tumor cells were quantified to determine tumor burden within the areas quantified. HNA-positive tumor cells were then assessed for double-labeling with or Ki67. To calculate proliferation index (the percentage of proliferating tumor cells for each animal), the total number of HNA-positive cells co-labeled with Ki67 across all areas quantified was divided by the total number of human nuclei-positive cells counted across all areas quantified. Differences in proliferation indices were calculated using unpaired, two-tailed Student’s *t*-tests (parametric data) or Mann-Whitney test (non-parametric data).

#### Statistical Analyses

Statistical tests were conducted using Prism (GraphPad) software unless otherwise indicated. Gaussian distribution was confirmed by the Shapiro-Wilk normality test. For parametric data, unpaired, two-tailed Student’s *t*-tests and one-way ANOVAs with Tukey post-hoc tests to further examine pairwise differences were used. For non-parametric data, Mann-Whitney test was used. The limiting dilution assay to test for neurosphere forming capacity was analyzed with a chi-squared test using the Extreme Limiting Dilution Analysis (ELDA) web-based tool (http://bioinf.wehi.edu.au/software/elda/). A level of *P* < 0.05 was used to designate significant differences. Based on the variance of xenograft growth in control mice, we used at least 3 mice per genotype to give 80% power to detect an effect size of 20% with a significance level of 0.05.

Statistical analyses for proteomic and RNA-seq data are described above in the respective sections.

#### Data Availability

RNA-seq data are deposited in GEO, accession number GSE99045

## Acknowledgements

The authors gratefully acknowledge support from the National Institute of Neurological Disorders and Stroke (NINDS R01NS092597 to M.M.), National Cancer Institute (1F31CA200273 to H.V.), Liwei Wang Research Fund (to M.M.), Department of Defense (NF140075 to M.M.), McKenna Claire Foundation (M.M.), Alex’s Lemonade Stand Foundation (M.M.), The Cure Starts Now Foundation and DIPG Collaborative (M.M.), Lyla Nsouli Foundation (M.M.), Unravel Pediatric Cancer (M.M.), California Institute for Regenerative Medicine (CIRM RN3-06510 to M.M.), Childhood Brain Tumor Foundation (M.M.), Matthew Larson Foundation (M.M.), V Foundation (M.M.), the Joey Fabus Childhood Cancer Foundation (M.M.), the Wayland Villars DIPG Foundation (M.M.), the Connor Johnson, Zoey Ganesh, Abigail Jensen, and Declan Gloster Memorial Funds (M.M.), N8 Foundation (M.M.), Virginia and D.K. Ludwig Fund for Cancer Research (M.M.), Child Health Research Institute at Stanford Anne T. and Robert M. Bass Endowed Faculty Scholarship in Pediatric Cancer and Blood Diseases (M.M.), NIH P50 CA168504 (J.J.Z), P50 CA165962 (J.J.Z.), R35 CA210057 (J.J.Z), and Breast Cancer Research Foundation (J.J.Z) and the intramural programs of the National Center for Advancing Translational Sciences and the National Cancer Institute (C.J.T.)

## Author Contributions

H.S.V., L.T.T., P.J.W, S.G., J.L, D.Y.D., P. J. M. conducted experiments. H.S.V. and M.M. contributed to experimental design. H.S.V, S.N., S.G., J.L., C.J.T. and M.M. contributed to data analysis. J.N and J.J.Z. developed and provided the breast cancer brain metastasis xenograft model. All authors contributed to manuscript editing. M.M. and H.S.V. wrote the manuscript. M.M. supervised all aspects of the work.

## Author Information

Correspondence and request for materials should be addressed to M.M. M.M. and H.S.V. declare that Stanford University has filed a patent application related to this work. RNA-seq data are deposited in GEO, accession number GSE99045

## Extended Data Figure Legends

**Extended Data Figure 1: Engraftment is equivalent in Nlgn3 knockout and wild type mice a,** *In vivo* bioluminescence imaging of SU-pcGBM2 xenografts two weeks following xenograft in WT;NSG (“WT”; left) or *Nlgn3* KO;NSG mice (“KO”; right). The heat map superimposed over the mouse heads represents the degree of photon emission by cells expressing firefly luciferase. **b**, Absolute flux of pHGG cells in identically manipulated WT;NSG (n=11) and *Nlgn3* KO;NSG (n=14) mice, measured by IVIS imaging two weeks post-xenograft illustrates no significant difference in tumor engraftment. Mann-Whitney test, n.s. = not significant (P > 0.05). Data are shown as mean +/- s.e.m.

**Extended Data Figure 2: Microenvironmental Nlgn3 is necessary for pediatric GBM growth.** Data from main Figure 1 shown on the same axis (**a**) and with each independent cohort color coded for comparison of littermates (**b**). Data illustrate growth of pHGG (SU-pcGBM2) xenografts in identically manipulated WT;NSG (black dots, n=11) and *Nlgn3*^y/-^;NSG (grey dots, n=14) mice, measured by IVIS imaging (fold change in total photon flux) and shown at 6, 12, 18, and 24 weeks post-xenograft. Data were replicated in five independent cohorts (litters) of mice xenografted with different cell preparations on different days and the data from these five biological replicates are shown combined with each cohort color-coded (i.e. littermates are shown in the same color). **P<0.01, ****P<0.0001, Mann-Whitney test. Data are shown as mean +/- s.e.m.

**Extended Data Figure 3: Nlgn3 is necessary for the full proliferative effect of brain slice conditioned media**

Schematic representation of active conditioned media generation (left). Proliferation index (EdU+ and DAPI co-positive nuclei/total DAPI+ nuclei) of pHGG cells (SU-pcGBM2) exposed to plain media (aCSF), optogenetically stimulated *Nlgn3* WT cortical slice CM, or optogenetically stimulated *Nlgn3* KO cortical slice CM (one-way ANOVA, *F* = 30.8, *P*<0.001). *P<0.05, ***P < 0.001

**Extended Data Figure 4: Neuroligin-1 does not stimulate glioma proliferation**

Proliferation index of patient-derived pediatric cortical glioblastoma (SU-pcGBM2) cells as measured by EdU incorporation 24 hrs after *in vitro* exposure to recombinant human neuroligin-1 (NLGN1) at concentrations ranging from 0-100 nM. n = 3 wells per condition. Data are presented as the mean EdU+/DAPI+ fraction +/- s.e.m. n.s. = not significant (p > 0.05) by one-way ANOVA.

**Extended Data Figure 5: Gene expression changes induced by neuroligin-3 in glioma**

**a**, Scatterplot showing SU-pcGBM2 (n=2) gene expression changes following 16 hours of treatment with vehicle (~1% DMSO) or NLGN3 (100 nM). The x-axis shows mean FPKM value in vehicle treated cells and the y-axis shows log2(fold-change) of NLGN3 over vehicle. Points shown in red represent genes showing statistically significant change (adjusted p value <0.1, Benjamini-Hochberg). **b**, Gene Ontology Biological Processes enriched in significantly upregulated genes with NLGN3 treatment, as identified by DAVID^49,50^ with p values shown with Benjamini-Hochberg adjustment. Genes associated with each GO BP term shown in (**c**).

**Extended Data Figure 6: Efficiency of Cre driver mice**

Recombination rate of inducible Cre driver models 7 days after treatment with tamoxifen (100mg/kg for 5 days) in *Rosa26*::tdTomato^lox-stop-lox^ reporter mice. **a,** To assess the neuron-specific *CamKIIa*:CreER Cre driver, recombination efficiency was quantified as percent of NeuN+ neurons that co-express tdTomato+ in the cortex of either *CamKIIa*:CreER- or *CamKIIa*:CreER^+^ mice 7 days following completion of tamoxifen administration. **b,** To assess the OPC-specific Cre driver *PDGFRa*:CreER, recombination efficiency was quantified as number PDGFRa+ OPCs that co-express tdTomato in the cortex of either *PDGFRa*:CreER**-** or *PDGFRa*:CreER+ mice. n = 3 mice per group. Data are shown as mean +/- s.e.m.

**Extended Data Figure 7: NLGN3 shedding from glioma cells is regulated by NLGN3 exposure and is mediated by ADAM10**

**a**, NLGN3 Western blot illustrating neuroligin-3 secreted into CM from optogenetically stimulated *Thy1::ChR2*; NSG cortical slices (ChR2 stim slice) or SU-DIPGXIII xenograft-bearing *Thy1::ChR2*; NSG cortical slices (ChR2 stim slice with xenograft). **b**, NLGN3 western blot illustrating neuroligin-3 secreted into CM from wild type brain slices (WT), WT brain slices bearing xenografts of adult GBM SU-GBM035 (WT + xeno), or from Nlgn3 knockout brains bearing SU-GBM035 xenografts (Nlgn3 KO + xeno) in the absence (left 3 lanes) or presence (right 3 lanes) of 200 nM ADAM10 inhibitor GI254023X (+ADAM10i). **c**, NLGN3 western blot illustrating glioma cell secretion of NLGN3 in vitro at baseline media conditions (aCSF), following exposure to recombinant NLGN3 with subsequent washing (NL3), at baseline media conditions in the presence of ADAM10 inhibitor GI254023X (aCSF + AD10i) or following NLNG3 exposure in the presence of ADAM10 inhibitor (NL3+AD10i). **d**, mRNA expression levels of *ADAM10* in primary tumor and cultures of DIPG by RNA-seq with values reported as FPKM^16,26^ (left) and in 493 individual adult glioblastoma samples from TCGA^27^ (right). Values are reported as robust multi-array averages (RMA; right).

**Extended Data Figure 8**

**a**, Spheroid invasion index of SU-DIPGVI cells exposed to ADAM10i (0-5μM) at 24-, 48- and 72-hours expressed as the diameter of the sphere of glioma cells relative to the initial diameter at time 0-hours. **b**, Extreme limiting dilution assay (ELDA) data presented in main Figure 6c re-plotted here as a log fraction plot with the slope of the solid line representing the log-active cell fraction and confidence intervals shown as dotted lines. SU-pcGBM2 cells treated with ADAM10 inhibitor GI254023X at 0.5 μM (black), 1 μM (red) or 2 μM (green), with vehicle (DMSO) control (royal blue) or no DMSO (cyan) and analyzed for neurosphere formation at two weeks.

**Extended Data Figure 9: Brain Penetration of XL-784**

Brain tissue and plasma levels of XL-784 at various time points following a single 50 mg/kg i.p. dose in NSG mice as assessed by liquid chromatography/tandem mass spectrometry (LC-MS/MS). n = 3 mice at each data point. Data are shown as mean +/- st.dev.

**Extended Data Table 1: Brain penetration of ADAM10 inhibitor GI254023X**

A single 100 mg/kg dose was delivered intra-peritoneally in NSG mice, and tissue samples collected 30-minutes later for analysis using liquid chromatography/tandem mass spectrometry (LC-MS/MS). Brain tissue concentrations show reasonable penetration across the blood brain barrier, achieving 2-4 μM concentration of drug at this time point.

